# Programmed cell death recruits macrophages into the developing mouse cochlea

**DOI:** 10.1101/2021.09.17.460339

**Authors:** Vikrant Borse, Tejbeer Kaur, Ashley Hinton, Kevin Ohlemiller, Mark E. Warchol

## Abstract

Programmed cell death (PCD) plays a critical role in the development and maturation of the cochlea. Significant remodeling occurs among cells of the greater epithelial ridge (GER) of Kölliker’s organ, leading to tissue regression and formation of the inner sulcus. In mice, this event normally occurs between postnatal days 5-15 (P5-15) and is regulated by thyroid hormone (T3). During this developmental time period, the cochlea also contains a large population of macrophages. Macrophages are frequently involved in the phagocytic clearance of dead cells, both during development and after injury, but the role of macrophages in the developing cochlea is unknown. This study examined the link between developmental cell death in the GER and the recruitment of macrophages into this region. Cell death in the basal GER begins at P5 and enhanced numbers of macrophages were observed at P7. This pattern of macrophage recruitment was unchanged in mice that were genetically deficient for CX3CR1, the receptor for fractalkine (a known macrophage chemoattractant). We found that injection of T3 at P0 and P1 caused GER cell death to begin at P3, and this premature PCD was accompanied by earlier recruitment of macrophages. We further found that depletion of macrophages from the developing cochlea (using CX3CR1^DTR/+^ mice and treatment with the CSF1R antagonist BLZ945) had no effects on the pattern of GER regression. Together, these findings suggest that macrophages are recruited into the GER region after initiation of developmental PCD, but that they are not essential for GER regression during cochlear remodeling.

## Introduction

The cochlea of the inner ear detects sound vibrations and transmits information to the auditory brainstem. Cochlear development is a complex process, and any defects in developmental patterning can lead to congenital hearing loss (**Korver et al., 2017, Anniko 1983, Kamilya et al., 2001, Rueda et al., 1987**). The developing cochlea also undergoes considerable cellular remodeling, in which certain cell populations die and are removed from surrounding tissues. In mice, such remodeling occurs during the first two postnatal weeks (**Anniko 1983, Kamilya et al., 2001, Rueda et al., 1987, Dayaratne et al., 2014, Ruben 1967, Sohmer and Freeman 1995**). Among the most critical remodeling events is the regression of a columnar epithelial structure known as the greater epithelial ridge (GER, also known as Kölliker’s organ). This event occurs via programmed cell death (PCD) (**Anniko 1983, Kamilya et al., 2001, Rueda et al., 1987, Dayaratne et al., 2014, Ruben 1967**), and creates a large cavity known as the inner spiral sulcus. This process is vital for the normal onset of hearing (**Dayaratne et al., 2014, Ruben 1967, Sohmer and Freeman 1995, Lukashkin et al., 2010**), and genetic deficits that prevent PCD result in profound hearing loss (**Makishima et al., 2011, Takashaki et al., 2001, Kuida et al.,1996, Morishita et al., 2001**). However, the signals that initiate PCD in the developing cochlea have not been completely identified.

Thyroid hormone signaling is critical for normal cochlear development **(Ng et al., 2013, Kelley and Forrest 2001)**. Hypothyroidism and mutations in the thyroid hormone receptor β gene (THRB) have been linked to hearing loss (**Kelley and Forrest 2001, Christ et al., 2004, Johnson et al., 2007, Mustapha et al., 2009, Sunderesan et al., 2015**). Notably, thyroid hormone has been shown to initiate regression of the GER during cochlear development (**Kelley and Forrest 2001, Peeters et al., 2015**). Mutations in THRB lead to delayed GER remodeling (**Kelley and Forrest 2001**), while ectopic treatment with T3 (3,5-triiodo-L-thyronine) at P0 and P1 induces premature GER remodeling (**Peeters et al., 2015**).

Programmed cell death also requires the clearance of dying cells. In many developing tissues, the removal of cellular debris is mediated by macrophages, which recognize apoptotic cells via pattern recognition molecules and phagocytic receptors (**Gordon and Plüddemann 2018, Wynn et al., 2013, Varol et al., 2015, Davies et al., 2013, Ergwin and Henson 2008, Henson and Hume 2006**). Resident macrophages are present in the developing and mature cochlea (**Warchol 2019, Hirose et al., 2017, Brown et al., 2017**), and deletion of macrophages results in hearing loss caused by excessive glial cells, abnormal myelin formation and edema in stria vascularis. Macrophage number transiently increases during cochlear development and declines as the cochlea matures (**Brown et al., 2017, Dong et al., 2019**), but the role of macrophages in cellular remodeling of the developing cochlea is not known. In this study, we examined the involvement of macrophages in the clearance of dying cells during GER remodeling. We found that macrophages were recruited into the GER after the initiation of cell death, and that those macrophages were engaged in the removal of cellular debris. In addition, we observed that early induction of PCD by ectopic injection of T3 hormone leads to earlier recruitment of macrophages in the GER region. Finally, we found that macrophage depletion did not affect GER regression in the developing mouse cochlea. Together, these findings suggest that developmental cell death leads to macrophage recruitment into the cochlea. Although such macrophages are actively involved in the clearance of cellular debris, they are not essential for GER regression.

## Results

### Recruitment of macrophages into the GER region of the developing mouse cochlea

The mouse cochlea undergoes considerable structural changes during the first two postnatal weeks **(Anniko 1983, Kamilya et al., 2001, Rueda et al., 1987, Dayaratne et al., 2014, Ruben 1967,).** One critical event is the death of columnar epithelial cells of the GER, leading to the opening of the inner sulcus. This process progresses along the tonotopic gradient of the cochlea, starting at the base (high frequency region) and then moving towards the apex (low frequency region) **(Anniko 1983, Kamilya et al., 2001, Rueda et al., 1987, Dayaratne et al., 2014, Ruben 1967,).** Numerous macrophages are also present in the developing cochlea (**Dong et al., 2019**), but the role of macrophages in cellular remodeling is unknown. To resolve this issue, we first characterized the patterns of programmed cell death and the corresponding recruitment of macrophages in the GER of developing cochlea, using CX3CR1^*GFP*/+^ mice, which express GFP in macrophages, microglia, monocytes and related cells (**Kaur et al., 2015, Diehl et al., 2013**). Specimens were fixed at various developmental timepoints and prepared as whole mounts or mid-modiolar frozen sections. Immunolabeling was used to enhance GFP fluorescence in macrophages, and to identify apoptotic cells (cleaved caspase-3), and neurons (TUJ1+NF). DAPI staining facilitated the identification of pyknotic nuclei (a hallmark of apoptosis). Increased numbers of dying cells with pyknotic nuclei and/or immunolabeled for activated caspase-3 were observed in the basal GER region between P5-10, with the number peaking at P7 (Fig. 1A-D’, F). Over the next 2-3 days, the appearance of pyknotic nuclei progressed from the base toward the apex, and the number of pyknotic nuclei in the apical GER peaked at P10 (Fig. 1C, F). Macrophages were also observed in the GER between P7-13. Large numbers of macrophages initially appeared in the basal GER and then spread toward the apex. The numbers of macrophages in the basal and apical GER regions peaked at P10 and P13, respectively (Fig. 1A-D’, G). The spatial distribution of macrophages followed a similar base-to-apex pattern that was observed for developmental cell death, with a temporal delay of ~3 days (Fig. 1A’). By P13, very few apoptotic cells were present within the GER region (Fig. 1D-D’), but macrophage numbers in the GER remained elevated after cell death had terminated (Fig. 1G). We also observed cleaved-caspase 3-labeled (i.e., apoptotic) cells being engulfed by macrophages (Fig. 1E). Similar patterns of cell death and macrophage recruitment were observed in mid-modiolar sections of the developing cochlea (Fig. 2A-E). Macrophages in the neonatal cochlea possessed both ameboid and ramified morphologies, while those in the mature cochlea were predominantly ramified (Fig. 2F-G). The neonatal cochlea also contained larger numbers of macrophages than were present in mature cochleae. Taken together, these observations indicate that macrophages are recruited into the cochlea in response to developmental cell death and are actively involved in clearing dying cells from the GER region.

**Figure 1:**
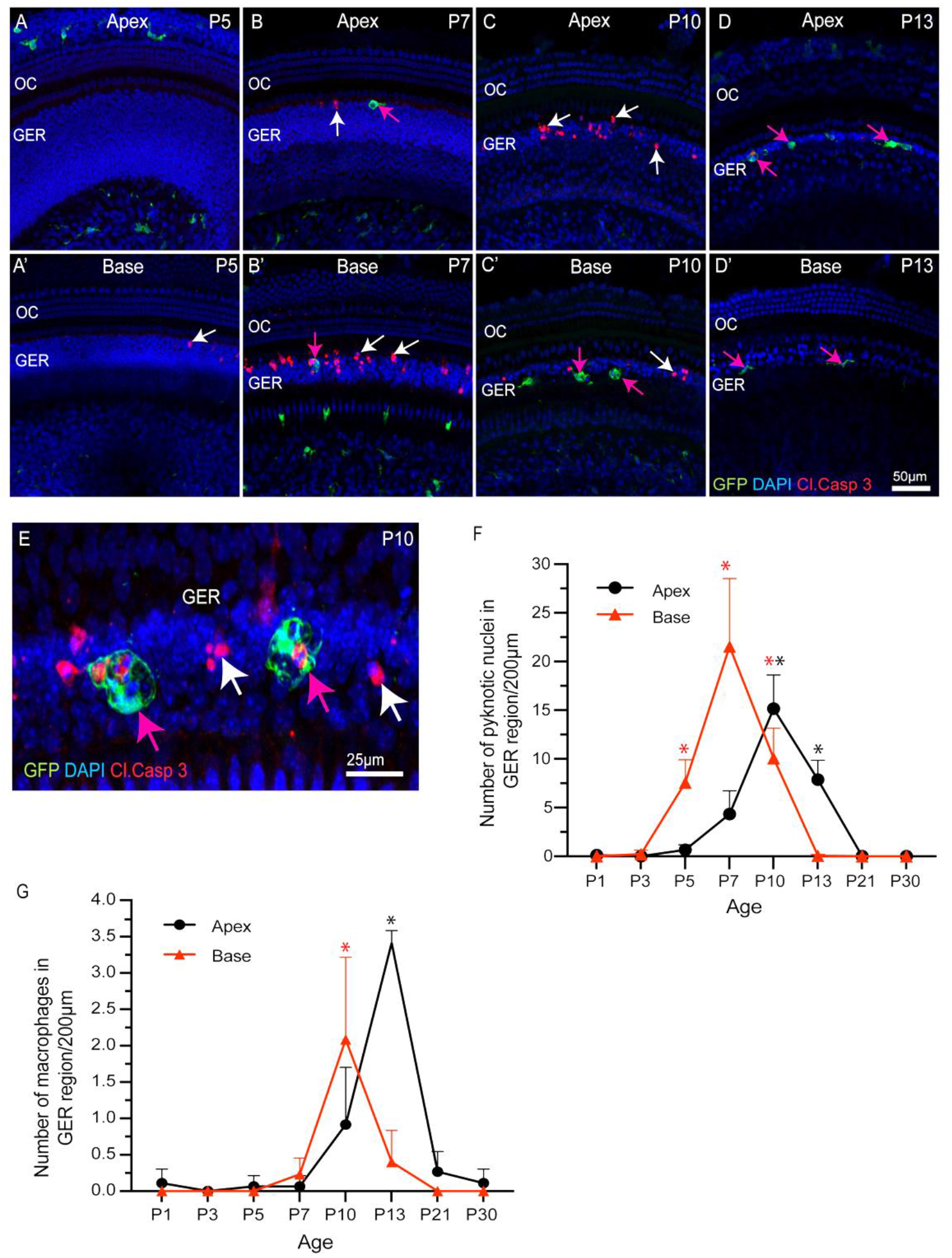
Progression of apoptosis and macrophage recruitment in the developing cochlea of CX3CR1^GFP/+^ mice. (A-D’) Sensory epithelium of the developing cochlea. Apoptotic cells were first observed in the basal region at P5 (white arrow, A’) and the apex at P7 (white arrow, B). Cell death continued at later time points (white arrows, B, C, B’, C’) and was accompanied by the recruitment of macrophages (magenta arrows). (E) A representative image of GFP-labeled macrophages engulfing pyknotic nuclei and activated caspase-3-labeled cells in the GER region at P10. (F) Number of pyknotic nuclei (a hallmark of apoptotic cells) in the GER region, as a function of developmental time. Cell death in the basal region peaked at P7, and in the apical region at P10. (G) Number of macrophages in the GER region, as a function of developmental time. The number of macrophages in both the basal and apical regions peaked at ~3 days after the time of maximal cell death. Labels: Red-cleaved caspase 3, Green-GFP+ macrophages, Blue-DAPI. White arrows: cleaved caspase-3-labeled (apoptotic) cells, magenta arrows: GFP+ macrophages. Abbreviations: - OC: organ of Corti, GER: Greater Epithelial Ridge, *p: p < 0.05, relative to P1. (N=3-6).

**Figure 2:**
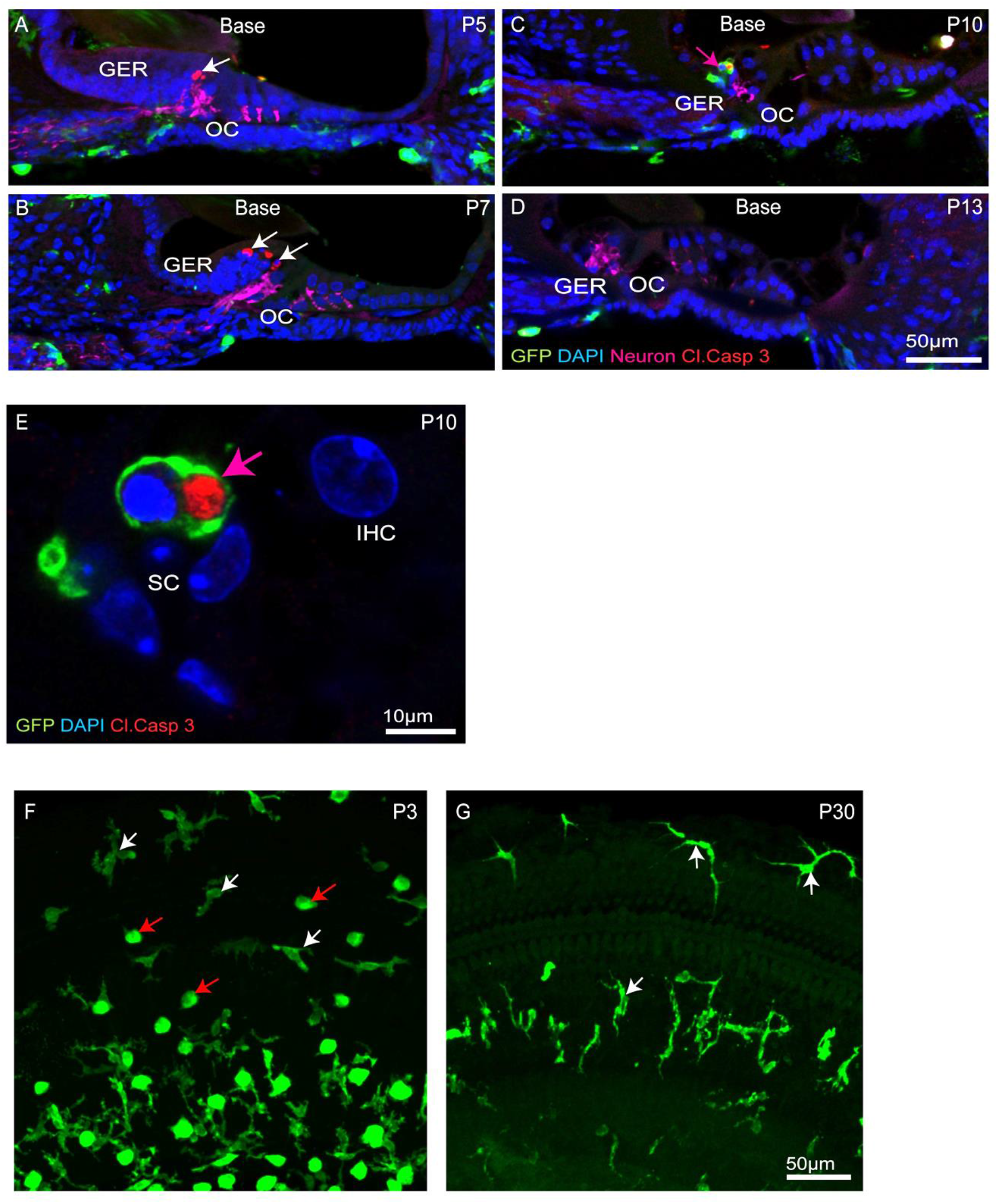
Progression of apoptosis and macrophage infiltration in the basal region of the developing cochlea. (A-D) Mid-modiolar sections of the developing cochlea. Apoptotic cells (white arrows, cleaved caspase-3, red) were observed at P5-10. Such cells were often engulfed by macrophages (magenta arrow, GFP, green) (E) Mid-modiolar section image showing GFP-labeled macrophage engulfing pyknotic nuclei at P10. (F-G) Presence of ameboid (red arrow) and ramified (white arrow) macrophages in cochlear whole mounts. Labels: Red-cleaved caspase-3, Green-GFP+ macrophages, Blue-DAPI. Abbreviations: - OC: organ of Corti, GER: Greater Epithelial Ridge, IHC: Inner Hair Cell, SC: Supporting Cell (N=3-6).

### Fractalkine signaling is not required for macrophage recruitment into the developing cochlea

Apoptotic cells secrete a number of signaling molecules that are chemoattractants for phagocytes (**Hirose et al., 2017, Park and Kim 2017, Sokolowski et al., 2014, Kaur et al., 2015, Diehl et al., 2013**). Examples of such ‘find-me’ signals include lysophosphatidylcholine (LPC), sphingosine-1-phosphate (S1P), nucleotides (ATP and UTP) and the CX3C motif chemokine ligand 1 (CX3CL1, also called as fractalkine) (**Park and Kim 2017**). CX3CL1 is the sole ligand for the CX3CR1 receptor, which is expressed by macrophages, microglia, monocytes and related cells. One function of fractalkine signaling is to meditate chemoattraction between dying cells and macrophages (**Park and Kim 2017, Sokolowski et al., 2014, Kaur et al., 2015**). To investigate the role of fractalkine signaling in macrophage recruitment and GER apoptosis, we compared macrophage numbers and cell death in the developing cochleae of CX3CR1^GFP/+^ mice (which possess one allele of *CX3CR1* and retain sensitivity to fractalkine) and CX3CR1^GFP/GFP^ mice (in which both copies of *CX3CR1* had been replaced with the gene for GFP). Cochleae were collected from mice of both genotypes at P5, P7, P10 and P13 and processed as whole mounts. Samples were immunolabeled for GFP (to enhance the fluorescent signal in macrophages), cleaved caspase-3 (to identify apoptotic cells), and stained with DAPI. These specimens displayed similar patterns of cell death and macrophage recruitment to those described above (Fig. 3). Apoptotic cells were first detected in the basal region of the GER at P5 (Fig. 3A’), and progressed towards the apex between P7-P10. By P13, few apoptotic cells were detected in the apical region and no apoptotic cells were found in the basal GER (Fig. 3A-D’, E). Macrophages were first apparent in the basal GER at P7 and their numbers moved apically at later ages (Fig. 3F). Macrophages remained present at P13, after cell death in the basal GER region had ended. The spatial and temporal patterns of macrophage recruitment were similar in both CX3CR1^*GFP/+*^ and CX3CR1^*GFF/GFP*^ (Fig. 1 and Fig. 3), and there was no statistically significant difference between the number of recruited macrophages and pyknotic nuclei in the GER region of CX3CR1^*GFP/+*^ vs. CX3CR1^*GFF/GFF*^ mice (Fig. 3E-F). These results indicate that fractalkine signaling is not essential for apoptosis or for recruitment of macrophages in the developing GER.

**Figure 3:**
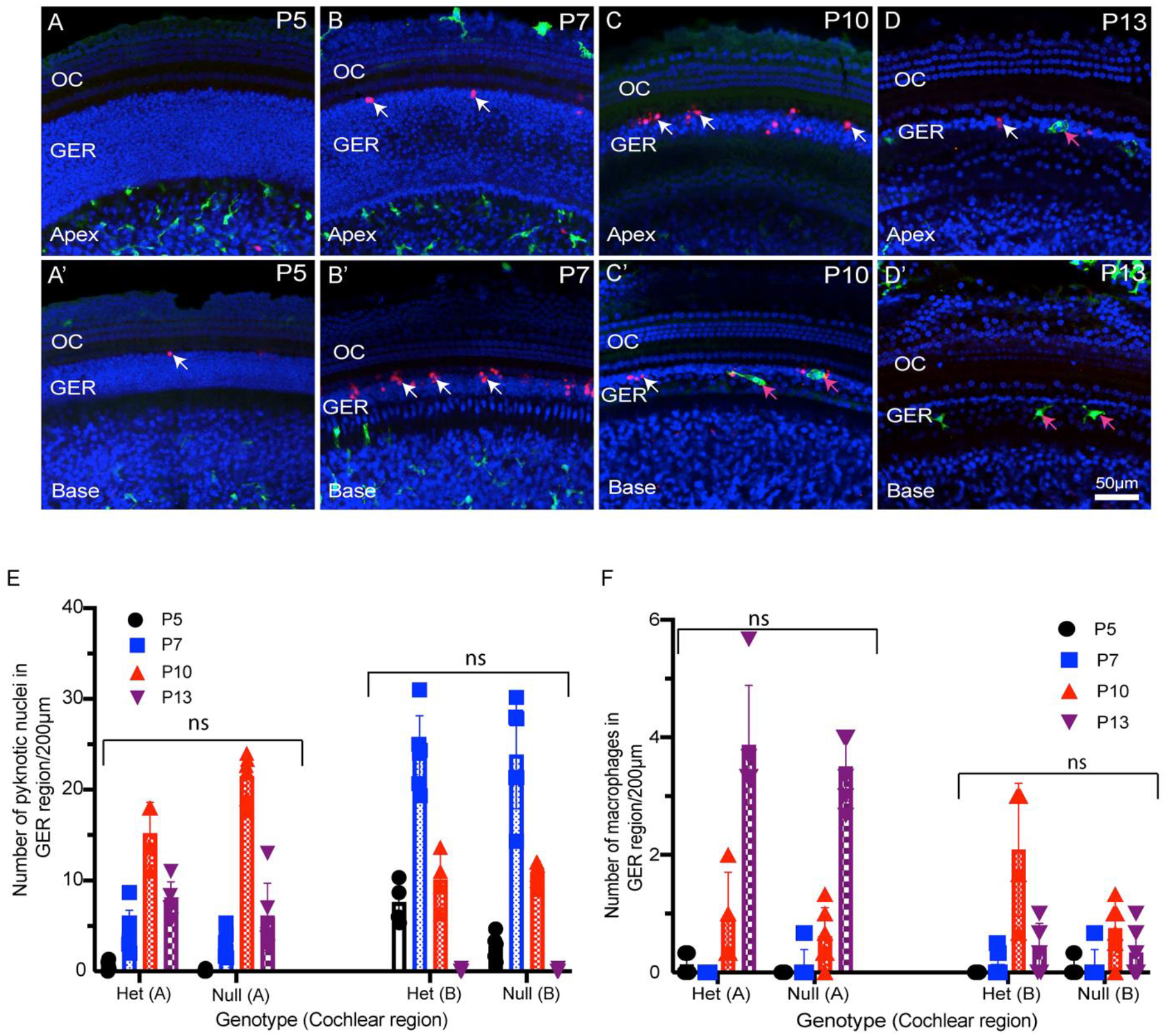
Progression of developmental apoptosis and macrophage recruitment is not affected by genetic deletion of CX3CR1. (A-D’) Sensory epithelium of the developing cochlea of CX3CR1-null mice. The patterns of cell death (white arrows, cleaved caspase-3) and macrophage recruitment (magenta arrows, GFP) are similar to those observed in WT mice (e.g., Fig. 1). (E) Total number of pyknotic nuclei in the GER region of CX3CR1-null mice vs CX3CR1^GFP/+^ mice. (F) Number of macrophages in the GER region of the developing cochlea of CX3CR1-null mice vs. CX3CR1^GFP/+^ mice. Labels: (A-D’) Red-cleaved caspase 3, Green-GFP+ macrophages, Blue-DAPI. White arrow-cleaved caspase-3-labeled cells, and magenta arrows-GFP+ macrophages. Abbreviations: - OC: organ of Corti, GER: Greater Epithelial Ridge, ns: not significant. (N=3-6).

### Thyroid hormone (T3) promotes premature cell death and early recruitment of macrophages

Thyroid hormone plays an important role in cochlear development and in the onset of hearing function **(Ng et al., 2013, Kelley and Forrest 2001)**. Both hypothyroidism and mutation of the thyroid hormone receptor β *(THRB)* gene delay the onset of GER remodeling and result in subsequent hearing loss (**Kelley and Forrest 2001, Christ et al., 2004, Johnson et al., 2007, Mustapha et al., 2009, Sunderesan et al., 2015**). It has also been shown that treating early neonatal mice with T3 hormone induces premature GER remodeling and accompanying hearing loss **(Peeters et al., 2015).** We investigated whether treatment with thyroid hormone (T3) also leads to earlier macrophage recruitment. Early apoptosis was induced in CX3CR1^*GFP*/+^ mice by injecting T3 at P0 and P1 (1.5 μg/pup, s.c.). We collected cochleae from both T3-treated and saline-injected (control) mice at various developmental time points. Whole mount specimens were labeled with anti-GFP (to enhance GFP signal in macrophages, cleaved caspase-3 (apoptotic marker), and DAPI. Both T3- and saline-treated cochleae showed a base-to-apex pattern of GER remodeling. However, treatment with T3 resulted in a significant increase in cell death at P3, which was followed by early recruitment of macrophages into the GER region at P5 (Fig. 4A). In contrast, cell death in saline treated samples began at P5 and was followed by recruitment of macrophages at P7 (Fig. 4A-E). Similarly, cell death in T3-treated cochleae terminated earlier than in control specimens (Fig. 4A-E). We also quantified hearing function in both T3- and saline-treated animals, using ABRs. Consistent with previous studies (**Peeters et al., 2015)**, we observed significantly elevated hearing thresholds in the T3-treated animals, when compared with saline-treated animals (Fig. 4F). Together, these data suggest that systemic injection of T3 hormone at P0 and P1 promotes premature apoptosis in the GER, which is followed by early recruitment of macrophages. This outcome is consistent with a causal link between developmental cell death and macrophage recruitment.

**Figure 4:**
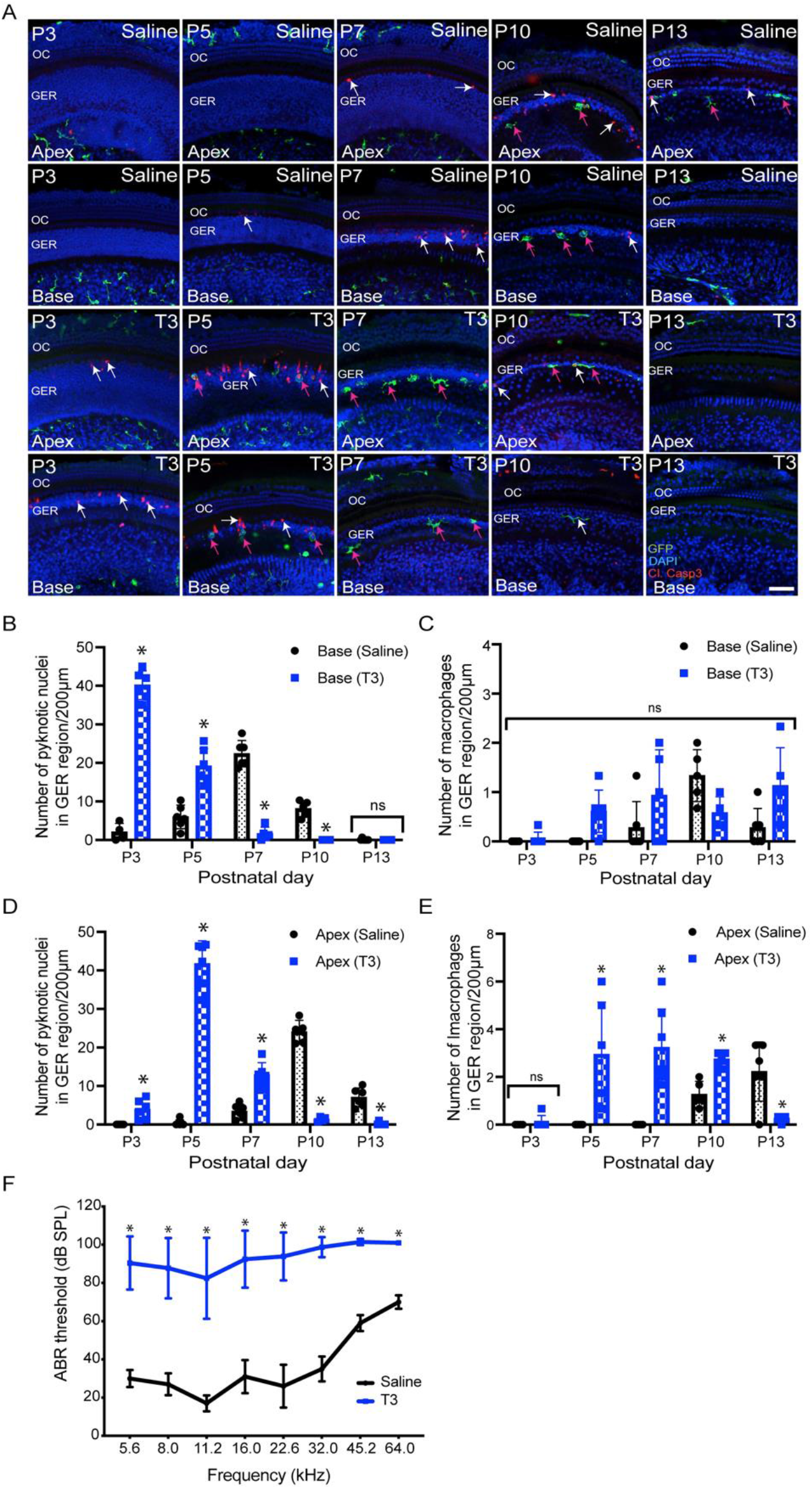
Thyroid hormone (T3) induces early apoptosis and macrophage infiltration in the GER region of the developing cochlea. (A) Sensory epithelia of developing cochleae in saline-treated (control) and T3-treated CX3CR1^GFP/+^ mice. Note that dying cells (white arrows) and macrophages (magenta arrows) are present ~2 days earlier in mice that received T3. (B, D) Number of pyknotic nuclei in the GER region of the developing cochlea of T3-treated and control (saline) mice. (C, E) Number of macrophages in the GER region of the developing cochlea of T3-treated and control mice. (F) ABR thresholds in T3-treated and saline treated control adult mice. ABR’s were assessed at P30. Treatment with T3 at P0-1 led to a ~60 dB elevation in ABR thresholds across the entire frequency range. Labels: Red-cleaved caspase-3, Green-GFP+ macrophages, Blue-DAPI. Abbreviations: - OC: organ of Corti, GER: Greater Epithelial Ridge, *p: p < 0.05, relative to saline treated, ns-not significant. (N=3-6).

### Deletion of macrophages does not affect GER remodeling

We investigated whether deletion of macrophages delays or impairs GER remodeling during cochlear development. We treated CX3CR1^DTR/+^ mice and wild type control mice with diphtheria toxin (**Diehl et al., 2013)** at P0, P2, P4, P6, P8, P10 and P12. Multiple doses of DT were delivered to prevent macrophage repopulation during the experimental period. Cochleae were fixed at P5, P7, P10 and P13 and processed as whole mounts (Fig. 6A). Samples were immunolabeled for CD45 (to identity macrophages) and cleaved caspase-3 (to identify apoptotic cells). Cell nuclei were also stained with DAPI. As depicted in Fig. 5B-D’, DT treatment led to nearly-complete deletion of macrophages from the developing cochlea of CX3CR1^DTR/+^ mice. Macrophage infiltration in the GER region of control animals was normal, but no macrophages were observed in the GER region of CX3CR1^DTR/+^ mice (Fig. 5E-L’). However, we observed normal patterns of apoptosis and GER regression in the developing cochlea of the CX3CR1^DTR/+^ and control mice (Fig. 5E-L’). Except in the apical region of P13 cochleae, we observed no significant increase in number of apoptotic cells in CX3CR1^DTR/+^ mice, in comparison with controls. These data suggest that depletion of macrophages may cause a slight delay in the clearance of cellular debris in the apex, but does not otherwise affect GER regression (Fig. 5M-N). Interestingly, we also noted that the DT treatment leads to loss of OHCs (Fig. 6A-B). We observed missing OHC nuclei and labeling for cleaved caspase-3 in both CX3CR1^DTR/+^ and WT control mice that were treated with DT at P10. No missing nuclei or apoptotic OHCs were observed in mice that did not receive DT.

**Figure 5:**
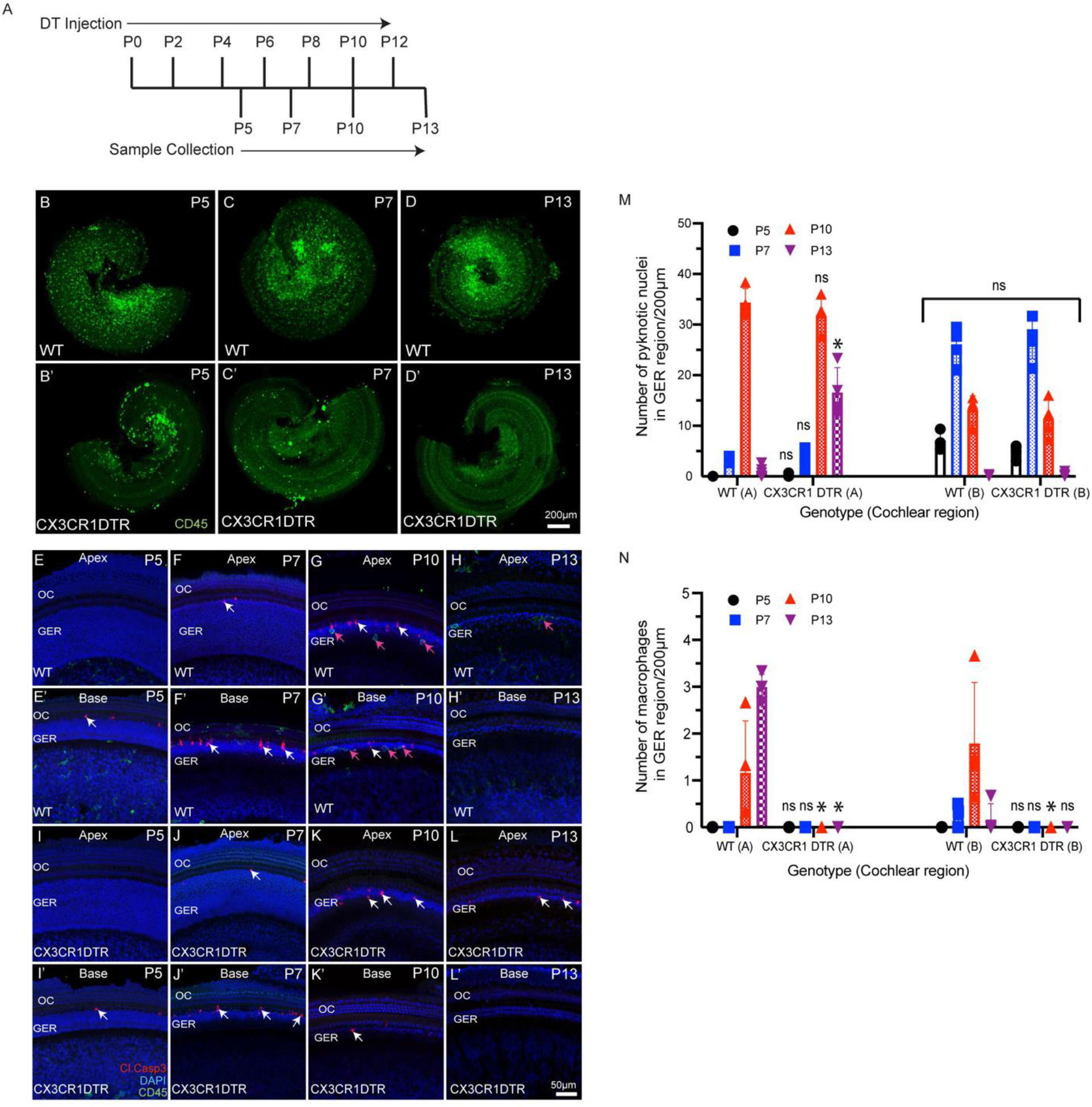
Depletion of macrophages in the cochlea of neonatal CX3CR-DTR mice. (A) Treatment and sample collection timeline. (B-D’) Cochlear whole mounts showing depletion of macrophages (Green: CD45) following DT treatment. (E-L’) Sensory epithelium of developing cochleae showing occurrence of cell death (red, cleaved caspase-3, white arrows) in normal and DT-treated cochleae. Although DT treatment eliminated macrophages in the cochleae of CX3CR-DTR mice, normal patterns of cell death were observed. (M) Number of pyknotic nuclei (blue: DAPI) in the GER region of developing cochleae. (N) Quantification of macrophages in the GER region of developing cochleae. Treatment with DT led to a large reduction in macrophage numbers. Abbreviations: OC: Organ of Corti, GER: Greater Epithelial Ridge, (*p: p value < 0.05, relative to WT, ns: not significant. (N=3-6).

**Figure 6:**
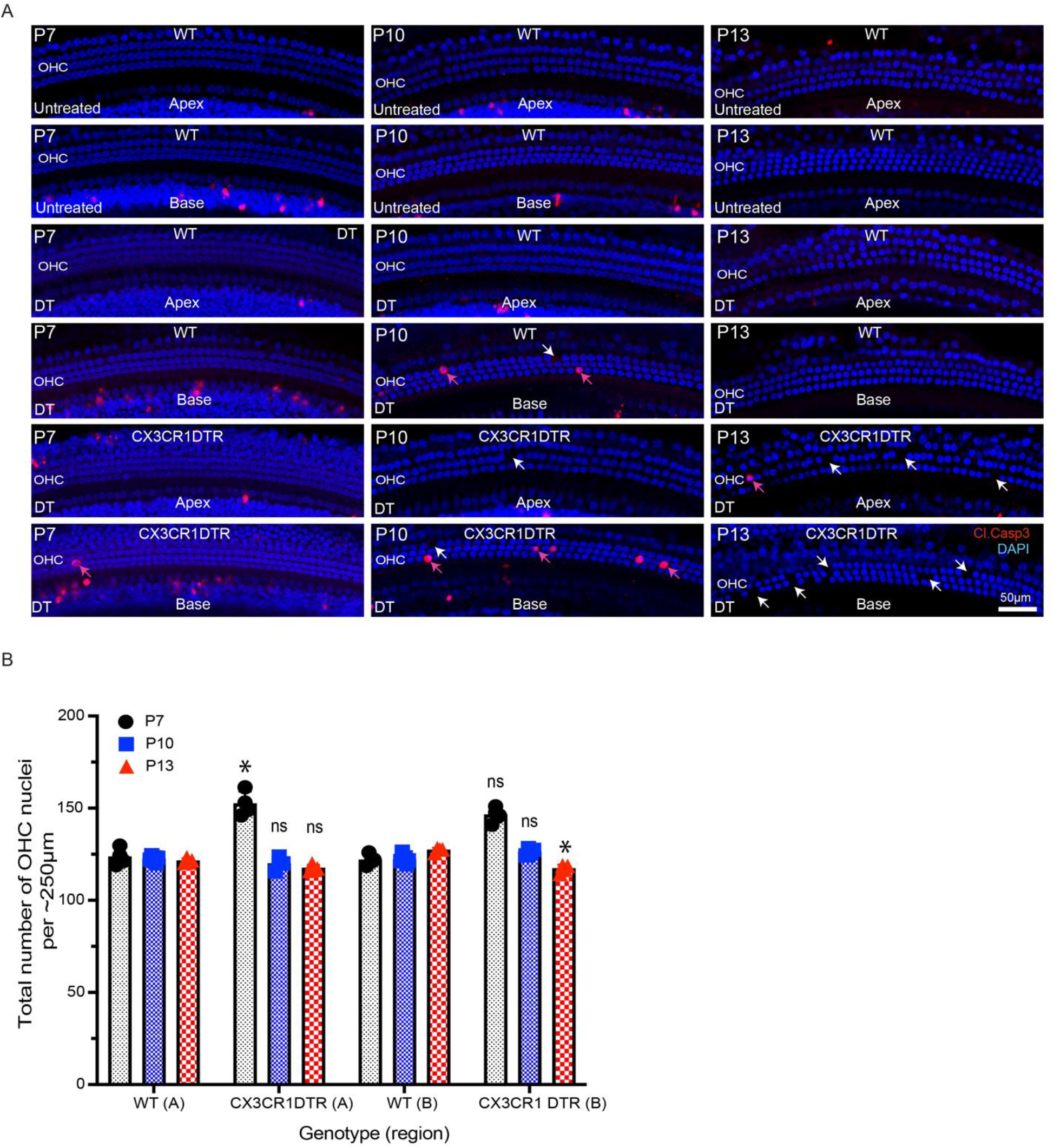
Treatment with DT leads to loss of OHCs from the developing cochlea. (A) Apoptotic cells (red, cleaved caspase-3) in the sensory epithelium of the developing cochlea. Treatment with DT causes death of a small number of OHCs, which is evident at P7-13. (B) Total number of OHCs in the cochlea of CX3CR-DTR and control mice. Abbreviations: - OHC: Outer hair cell, DT: Diphtheria toxin. *p: p < 0.05, relative to WT), ns: not significant. (N=3-6).

In addition to the CX3CR1^DTR/+^ mouse model, we also used a pharmacological approach to eliminate macrophages from the developing cochlea. CX3CR1^GFP/+^ mice were given s.c. injections of the CSF1R antagonist BLZ945 **(Milinkeviciute et al., 2019)** at P2, P4, P6, P8, P10 and P12. (Multiple doses of BLZ945 were delivered, in order to avoid macrophage repopulation.) Cochleae were fixed at P5, P7, P10 and P13, and processed as whole mounts (Fig. 7A). Treatment with BLZ945 eliminated nearly all macrophages from the developing cochlea (Fig. 7B-D’), including the GER region (Fig.7E-L’). Notably, we observed normal patterns of apoptosis and GER regression in the developing cochleae of both the BLZ945-treated and control animals (Fig.7M). Macrophage depletion led to no significant difference in the numbers of pyknotic nuclei in the GER, when compared to controls (Fig. 7M-N). Collectively, these findings suggest that macrophage depletion has no substantial effect on GER remodeling during development.

**Figure 7:**
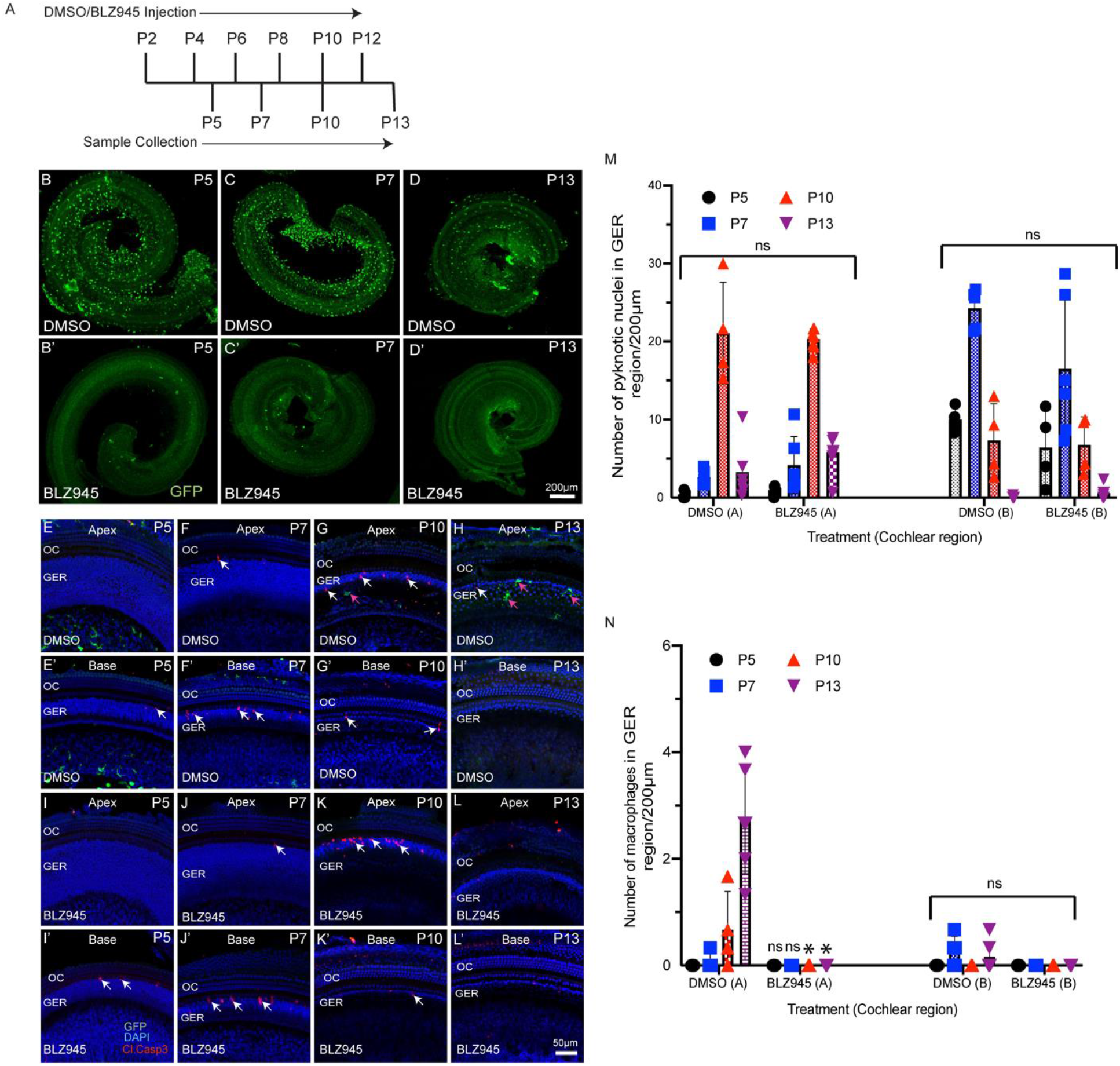
BLZ945 treatment eliminates macrophages in the developing cochlea. (A) Treatment and sample collection timeline. (B-D’) Cochlear whole mounts showing depletion of macrophages following treatment with the CSF1R antagonist BLZ945 (Green-GFP+ macrophages). (E-L’) Sensory epithelium of the developing cochlea showing apoptotic cells (red-cleaved caspase-3) and macrophages (green-GFP). Normal patterns of dying cells (white arrows) were observed in BLZ945-treated cochleae. (M) Number of macrophages in the GER region of the developing cochlea. Note that BLZ945 sharply reduces the number of macrophages in the GER. (N) Number of pyknotic nuclei in the GER region of the developing cochlea. Abbreviations: - OC: Organ of Corti, GER: Greater Epithelial Ridge, P: Postnatal Day. (*p: p < 0.05, relative to controls (DMSO), ns: not significant. (N=3-6).

## Discussion

This study was undertaken to examine the role of macrophages in cochlear development. We found that the spatial and temporal patterns of macrophage recruitment into the GER region of the developing cochlea is correlated with the pattern of programmed cell death in that region, i.e. starting from high frequency basal region and then moving towards low frequency apical region. We first observed apoptosis and macrophage recruitment in the GER region at P5 and P7, respectively (Fig. 1). In previous studies of cochlear development, apoptosis was first reported to occur at P7 (**Kamiya et al., 2001, Peeters et al., 2015**). This difference in timing may be due to differences in technique and mouse strain. We also observed macrophages engulfing apoptotic cells and pyknotic nuclei in the GER region and some macrophages that remained in the GER region after the wave of apoptosis had terminated. Our findings demonstrate that macrophages are recruited into the GER region about two days after apoptosis has begun. We also observed two morphologically-distinct populations of macrophages in the developing cochlea (ameboid and ramified), but only ramified macrophages were present in the mature cochlea. The significance of these morphological types is not clear. Microglia (the resident macrophages of CNS) also possess similar morphologies. Microglia with a ramified morphology are thought to be in a ‘resting’ or ‘quiescent’ state, while ameboid microglia are associated with tissue injury **(Savage et al., 2019).** However, the possible differential roles of ameboid vs. ramified macrophages during cochlear development are not understood.

Apoptotic cells release signaling molecules, known as ‘find-me’ signals, to promote the chemotaxis of macrophages to sites of cell death (**Park and Kim 2017**). A number of such signaling molecules have been identified. Fractalkine (CX3CL1) is a chemokine that can recruit macrophages to dying cells by activating the CX3CR1 receptor on macrophages (**Park and Kim 2017, Sokolowski et al., 2014, Kaur et al., 2015**). Previously, our lab has shown that disruption of fractalkine (CX3CL1/CX3CR1) signaling leads to reduced macrophage recruitment into sensory epithelium and spiral ganglion of the cochlea after selective hair cell ablation (**Kaur et al., 2015**). However, it was not known whether fractalkine also influences macrophage behavior during cochlear development. We investigated the role of fractalkine signaling by quantifying macrophage recruitment into the developing cochleae of CX3CR1-null (CX3CR1^GFP/GFP^) mice. We observed similar patterns of apoptosis and macrophage recruitment in both CX3CR1^GFP/GFP^ and CX3CR1^GFP/+^ (control) mice (Fig. 1 and Fig. 3), suggesting that fractalkine signaling is not required for normal macrophage recruitment into the developing cochlea. This finding suggests that other ‘find-me’ signals, such as LPC, S1P and/or nucleotides (ATP and UTP) might recruit macrophages into the neonatal cochlea (**Park and Kim 2017,**). It is notable that spontaneous release of ATP occurs during cochlear development (**Dayaratne et al., 2014, Park and Kim 2017, Wang et al., 2015**), and this may be one signal that recruits macrophages into the GER.

In order to determine the relationship between GER cell death and macrophage recruitment, we studied whether early induction of apoptosis by T3 hormone affected the recruitment of macrophages into the GER. Similar to Peeters et. al. (**Peeters et al., 2015**), we observed robust early apoptosis in the GER region of T3 treated animals. In addition, we noted that induction of premature apoptosis by T3 injection led to early recruitment of macrophages. Treatment with T3 caused cell death in the GER to begin approximately 4-5 days earlier than in normal cochleae and, in accordance with earlier studies, led to profound hearing loss (**Peeters et al., 2015**). We further found that T3 treatment led to correspondingly earlier macrophage recruitment into the GER, but that the delay between the onset of cell death and the appearance of macrophages was ~2 days in both normal and T3-treated cochleae. These observations support a causal relationship between cell death and macrophage recruitment, with dying cells releasing a chemoattractant that causes macrophages to migrate into the GER within ~2 days.

A prior study reported that short-term depletion of macrophages during cochlear development leads to hearing impairment, which was attributed to the presence of excessive glial cells, abnormal myelin formation, and defects in the stria vascularis (**Brown et al., 2017**). The present study employed two methods for depletion of macrophages: treating CX3CR1^DTR/+^ mice with DT (**Diehl et al., 2013**), and treating CX3CR1^GFP/+^ mice with the CSF1R antagonist BLZ945 (**Milinkeviciute et al., 2019**). Similar to earlier studies, we observed that multiple doses of DT and BLZ945 resulted in sustained near-complete deletion of macrophages from the developing cochlea (**Diehl et al., 2013, Milinkeviciute et al., 2019**). We further found that eliminating macrophages did not disrupt the normal pattern of apoptosis in the GER region of the developing cochlea. These results suggest that the cells in the GER region may engage in self-clearance by autophagy and/or by phagocytosis (**Monzack et al., 2015, Hou et al., 2019**). Similarly, in the mature ear, both macrophages and supporting cells have been shown to contribute to the phagocytic clearance of dead hair cells (**Monzack et al., 2015**). Recruited macrophages may assist GER cells in rapid clearance of dead cells, but they do not appear to be essential for GER regression during normal cochlear development.

In conclusion, the results of this study demonstrate that programmed cell death recruits macrophages in the GER of the developing cochlea and that macrophages engage in the phagocytic clearance of dead cells. However, elimination of macrophages does not affect the patterns of cell death and remodeling in the GER. Thus, although the developing cochlea contains a higher density of macrophages than are present in the mature cochlea, macrophages may not play an essential role in the development of the organ of Corti.

## Materials and Methods

### Animals

Studies used CX3CR1-GFP and CX3CR1-DTR mice (**Diehl et al., 2013**), of both sexes, on a C57BL/6 (B6) background. Samples were obtained at post-natal (P) day 1, P3, P5 P7, P10, P13, P21 and P30, in order to study the role of macrophages in cochlear development and function Each experimental group consisted of 3-6 animals, taken from multiple litters. CX3CR1-GFP mice express enhanced green fluorescent protein (EGFP) under control of the endogenous *CX3CR1* promoter (**Jung et al., 2000**). We generated CX3CR1^GFP/+^ mice by breeding CX3CR1-null (CX3CR1^GFP/GFP^) mice with wild type C57BL/6 mice. Similarly, we generated CX3CR1^DTR/+^ mice by breeding CX3CR1-null (CX3CR1^DTR/DTR^) mice with wild type C57BL/6 mice. All mice were housed in the animal facilities at Washington University, School of Medicine, and were maintained on a 12-hr/day-night light cycle with open access to food and water. All experimental protocols involving animals were approved by the Animal Studies Committee of the Washington University School of Medicine, in Saint Louis, MO.

### Treatment

At P0 and P1, mouse pups were subcutaneously injected with 1.5 μg of T3 (catalog# T6397, Sigma) (10 μl) or saline (control), in order to induce premature apoptosis and GER remodeling (**Peeters et al., 2015**). Saline injected animals were used as controls. Temporal bones from these mice were collected between P3 to P30. Also, some mice received subcutaneous injections of Diphtheria toxin (DT, Sigma), or the colony stimulating factor receptor 1 (CSF1R) inhibitor BLZ945 (MW: 398.48, MedChem Express HY-12768/CS-3971), to eliminate macrophages (**Diehl et al., 2013, Milinkeviciute et al., 2019**). In these studies, a single dose of 5 ng/gm DT was injected at P0, P2, P4, P6, P8, P10 and P12 in CX3CR1-DTR mice. Other mice received a single dose of 200 mg/kg BLZ945 at P2, P4, P6, P8, P10 and P12. Control mice received either saline or DMSO. Mice were euthanized and temporal bones were collected between P3 to P13.

### Genotyping

Tail-clip samples were obtained from CX3CR1-GFP (**Jung et al., 2000)** and CX3CR1-DTR mice (**Diehl et al., 2013**), and genotyping was performed by Transnetyx.

#### Histological methods

Animals were euthanized either by quick decapitation or deep anesthesia (Fatal Plus, 50mg/kg). Temporal bones were isolated and fixed overnight with 4% paraformaldehyde (in PBS) at 4° C. After fixation, cochleae were isolated and washed 3x for 5 min each in PBS, pH 7.4 at room temperature. All specimens, except those collected at P5, were then incubated for ~24 hr in 10% EDTA in PBS, pH 7.4 at room temperature. Specimens were thoroughly washed 3x for 5 min each in PBS, pH 7.4 at room temperature and were then dissected into whole mount samples. Other samples were prepared as frozen mid-modiolar sections (30μm thickness). Those cochleae were incubated in 10% EDTA in PBS, pH 7.4 at room temperature for 3-5 days and then washed 3x for 5 min each in PBS, pH 7.4 at room temperature. Cochlear whole mounts and mid-modiolar sections were immunolabel for GFP-expressing macrophages (rabbit anti-GFP antibody, catalog #A11122, Invitrogen, 1:500 or chicken anti-GFP antibody, catalog #1010, Aves, 1:500), neurons were labeled with mouse monoclonal anti-β-III tubulin (catalog #MMS-435P, Covance, 1:500), combined with mouse anti-Neurofilament (Catalog #2H3, Developmental Studies Hybridoma Bank, 1:100) antibodies, and apoptotic cells were labeled using rabbit anti-cleaved caspase-3 antibody (catalog # 9661, Cell signaling, 1:100). To avoid non-specific binding of the antibodies, samples were preincubated in a blocking solution consisting of 5% normal horse serum/0.2% Triton X-100 in PBS for 1 hr at room temperature. Samples were then incubated overnight with primary antibodies prepared in PBS with 2% normal horse serum and 0.2% Triton X-100 at room temperature. Samples were washed 3x for 5 min each in PBS, pH 7.4 at room temperature and then labeled with secondary antibodies, conjugated with Alexa-488, Alexa-568 or 555, and Alexa 647, (Life Technologies,1:500). Secondaries were prepared in PBS with 2% normal horse serum and 0.2% Triton X-100. Samples were incubated in secondaries for 2-3 hr at room temperature. The secondary solution also contained DAPI, in order to counterstain cell nuclei (catalog #D9542, Sigma-Aldrich, 1 μg/ml). All samples were washed 3x for 5 min each in PBS, pH 7.4 at room temperature, mounted on glass slides in glycerol: PBS (9:1), and coverslipped.

### Cellular imaging and analyses

Fluorescence images were obtained using an LSM 700 confocal microscope (Zeiss). For cochlear whole mounts and mid modiolar sections, Z-series images were obtained using the following objective lenses: 5x (~40-micron z-step-size), 20x (1 or 2 -micron z-step-size), or 63x (0.5 or 1.0-micron z-step-size). Images were processed and analyzed using Volocity 3D image analysis software (version 6.3, PerkinElmer) and Fiji (ImageJ2.0) (National Institutes of Health) and Adobe illustrator CS5.1.

### Auditory brainstem responses (ABR)

Hearing was assessed from one-month old mice using auditory brainstem responses. Mice were anesthetized using intraperitoneal injections of ketamine (100 mg/kg) and xylazine (20 mg/kg). Responses were recorded from subcutaneous electrodes placed at the vertex (active electrode), and behind the right ear (reference electrode). A ground electrode was placed on the back. ABR thresholds were measured in response to tone pips at 5.6, 8, 11.2, 16, 22.6, 32, 45.2, and 64 kHz, using 5ms tone pips (including 0.5 ms cosine^2^ rise/fall) at a repetition rate of 20 per second. The responses were amplified 10,000x, filtered (100 Hz to 3 kHz), and averaged using BioSig software (System 3; Tucker-Davis Technologies). Stimuli were presented in 5 dB steps from 5 dB to 100 dB SPL, in decreasing order, to the right ear. At each level, up to 1000 responses were averaged. The lowest sound level at which a recognizable and reproducible wave 1 and 2 complex was noted, and considered threshold. If hearing threshold was not present at 100 dB SPL (the typical maximum sound level in our system), the threshold value was assigned as 100 dB for graphing and statistical purposes.

### Macrophage, pyknotic nuclei and hair cells counts

Quantification was performed from 20X images, using Fiji software (ImageJ2.0) (National Institutes of Health). For counts of macrophages, hair cells and pyknotic nuclei, 2-3 areas from the apical and basal GER regions were selected. The Cell Counter plug-in was used for all counts. Macrophages were identified by strong GFP and CD45 labeling. Pyknotic nuclei were identified by small size and intense DAPI nuclear labeling, and apoptosis was verified by immunolabeling for cleaved caspase-3. All macrophages, pyknotic nuclei and hair cells counts were obtained manually, within the designed area of complete z-stack using Fiji software.

### Statistical analysis

All data analysis and statistical tests were carried out using GraphPad Prism version 6.0d. Data are presented as mean ± SD. Student’s t-tests or analyses of variance (ANOVA), followed by post hoc tests, were applied as appropriate. Results were considered statistically significant when p < 0.05.

## Acknowledgment

We thank Dr. Keiko Hirose (Washington University) for providing CX3CR1^DTR^ mouse line.

## Notes

**Funding**, Supported by grant R01DC015790 from the NIH (Mark E. Warchol)

**Conflict of Interest**, All authors state no competing financial interest.

### Competing Interest Statement

The authors have declared no competing interest.

